# Generative design and construction of functional plasmids with a DNA language model

**DOI:** 10.64898/2025.12.06.692736

**Authors:** Angus G. Cunningham, Linda Dekker, Anastasiia Shcherbakova, Chris P. Barnes

## Abstract

DNA language models offer a new paradigm for sequence design, yet their ability to generate functional genomic sequences remains underexplored. Plasmids act as a good testbed for evaluating DNA language model generation potential due to their simplicity and ease of construction. Here, we develop an end-to-end pipeline for generative design of *Escherichia coli* plasmid backbones, from large-scale data curation through fine-tuning, sampling, bioinformatic assessment, and candidate selection. A curated plasmid library was assembled from PlasmidScope and Addgene, and PlasmidGPT, a GPT-2-style DNA model, was fine-tuned on these corpora using circular-aware batching and random crops. Generations (1,000 per model) were produced under two prompting strategies: a minimal ATG seed to expose default tendencies, and a GFP cassette to enforce functional context. From 1000 generated synthetic plasmids, 16 candidates survived strict filtering and these were prioritised for wet-lab validation. Three shortlisted plasmids were synthesised and found to be functional, supporting growth, antibiotic resistance, and GFP expression in *E. coli*. These represent, to our knowledge, the first full AI-generated plasmids to be synthesised and validated *in vivo*. This work demonstrates that curated fine-tuning and prompt-aware generation enable DNA language models to progress from raw sequence sampling to experimentally testable plasmid designs. The approach offers a foundation for extending DNA design optimisation beyond *E. coli*, toward broader applications across engineering biology.

## Introduction

Plasmids are small, typically circular DNA molecules that replicate independently of the bacterial chromosome and form a foundation of modern synthetic biology [1]. Unlike the chromosome, which encodes the full complement of cellular functions, a plasmid usually carries only a few genes but is maintained autonomously within the host cell. This autonomy makes plasmids versatile vectors for introducing new traits into microbes; they underpin modern synthetic biology as compact, modular vehicles for gene delivery, expression, and circuit prototyping in bacteria. Designing a robust plasmid backbone, however, remains an iterative process: choices about replication origin, selection marker, backbone architecture, and overall size are interdependent and strongly influence copy number, stability, and expression [2].

Standard practice integrates sequence repositories such as Addgene [3], which provide extensive libraries of engineered plasmids, with computational tools that annotate and verify sequence features. Platforms like *pLannotate* identify origins, markers, and cloning sites [4], while similarity searches using *BLAST* confirm novelty and flag unwanted elements. Classification tools such as *MOB-typer* provide replicon and mobility typing [5], and user-facing software such as *SnapGene* enables visual design and cloning planning [6]. Registries like *SynBioHub* further support design sharing and versioning in standardised formats [7]. Despite these advances, plasmid design still relies heavily on expert-driven iteration and experimental validation: constructs must ultimately be synthesised or cloned, transformed into host strains, and tested for replication, selection, and expression outputs.

Engineered plasmids generally contain three core modules. The origin of replication (ORI) is the defining structural feature - a DNA sequence that recruits host replication machinery and determines both copy number and host range. High-copy origins in *E. coli* (e.g. ColE1 derivatives) can yield dozens of plasmids per cell, boosting gene dosage, while low-copy origins provide stable maintenance with less metabolic burden [2]. ORI choice therefore governs propagation and stability and remains a central design parameter. The second module is the selectable marker, most often an antimicrobial resistance gene (ARG), which ensures plasmid retention under antibiotic pressure. In common laboratory plasmids, such markers include *blaTEM* (ampicillin), *aph*(3) (kanamycin), or *catA* (chloramphenicol), chosen according to host background, compatibility, and biosafety constraints. Finally, plasmids carry genes of interest or reporters under regulatory control. Reporters such as green fluorescent protein (GFP) provide measurable outputs linked to gene expression, enabling rapid functional screening [8]. Coupled with promoters, ribosome binding sites, and terminators, these components make plasmids modular devices in which genetic parts can be swapped or recombined as needed. Modular assembly systems such as MoClo (Modular Cloning) and Golden Gate standardisation frameworks use type IIS restriction enzymes and defined part hierarchies to facilitate the exchange of promoters, ORFs, and regulatory elements within standardised backbones [9]. The Standard European Vector Architecture (SEVA) plasmid toolkit extends this modular paradigm by providing a unified syntax and assembly framework for bacterial vectors across species [10]. SEVA defines interchangeable modules for replication origins, antibiotic markers, and cargo regions, allowing systematic swapping of functional parts within a stable backbone format. Together with Golden Gate and MoClo systems, SEVA exemplifies the growing move toward standardized, interoperable plasmid design in synthetic biology. At the same time, large-scale sequencing of plasmids has revealed that plasmid DNA exhibits a characteristic “grammar”—regularities in composition and motif usage shared across diverse backbones [11].

Recent advances in artificial intelligence have enabled DNA sequences to be treated as biological text, giving rise to DNA language models (DNA LMs). These models, typically based on Transformer architectures [12], learn the statistical “grammar” of DNA by training on large corpora in a self-supervised manner. Motifs such as promoters or origins of replication act like words with meaning, while long-range dependencies (e.g. operons or regulatory networks) form higher-order context. By capturing these patterns, DNA LMs provide a foundation for both predictive tasks (e.g. promoter classification) and generative design (e.g. proposing new plasmid backbones). The first wave of DNA LMs adapted BERT-style masked language modelling to genomic data. DNABERT tokenised sequences into overlapping *k*-mers, learning motif representations useful for tasks such as promoter or splice site detection [13]. Its successor, DNABERT-2, replaced fixed *k*-mers with data-driven subwords via byte-pair encoding (BPE), reducing redundancy and capturing motifs of variable length [14]. Together with improved positional encodings (e.g. ALiBi), this enabled learning from much longer contexts. Similar approaches such as the Nucleotide Transformer adopted rotary embeddings (RoPE) to scale to large genomes, providing transferable nucleotide embeddings [15]. While primarily encoders rather than generators, these models established that Transformers can extract biologically meaningful features from raw sequence. Subsequent work shifted towards autoregressive generation—predicting nucleotides sequentially—to model entire genes or genomes e.g. DNAGPT, MegaDNA illustrate that genome or virus scale generation is feasible, with plasmid-scale sequences falling well within reach [16, 17]. Evo and its successor Evo 2 exemplify this direction, combining efficient attention mechanisms using the convolution-inspired operators with massive genomic corpora to achieve context windows up to megabases [18, 19, 20]. These models demonstrated that biological dependencies can be captured across whole genomes and recently found to create valid bacteriophages with fine-tuning [21]. Recently, models have been trained directly on engineered plasmid corpora. PlasmidGPT, a GPT-2–style decoder trained on ∼150,000 Addgene plasmids, internalised the common grammar of vector construction—origins, antibiotic markers, cassettes, and cloning sites—and can generate novel but realistic plasmids [22]. Its embeddings are also predictive of plasmid function, including host range and ORI family. OriGen focuses more narrowly on origins of replication, using generative modelling to propose novel replicons, some of which have already been shown experimentally to function [23]. These plasmid-specialised models are most directly relevant to backbone optimisation, as they couple generative capacity with domain-specific priors.

Building on these advances, this study develops an end-to-end framework for plasmid backbone generation and evaluation using DNA language models. First, novel *E. coli*–specific datasets were curated by combining PlasmidScope with 10,304 Addgene plasmids and filtered to yield two fine-tuning corpora (ft35k and ft15k) of increasing biological stringency [24, 3]. Second, a decoder-only DNA LLM (PlasmidGPT) was fine-tuned with tokenisation and collator choices that respect circularity and variable length [22]. Third, controlled sampling generated 1,000 sequences per model under two prompts: a weak ATG codon seed and a GFP-cassette seed anchoring generation around a promoter–RBS–GFP–terminator context. Sampling employed a base-pair budget stopping rule to target realistic lengths and avoided EOS tokens to maintain circular closure. Finally, an evaluation pipeline combined Prodigal and pLannotate for gene-level inspection [25, 4], BLAST-based ORI typing [26], AMRFinderPlus for antimicrobial resistance detection [27], repeat screening, and complementary similarity measures. A two-tier filtering scheme converted raw generations into candidates: broad realism checks (single ORI, intact ARGs, low repeat burden) followed by strict selection for synthesis-ready backbones that balance integrity, stability, and novelty. Finally, we synthesised and constructed the first examples of AI-generated plasmids expressing GFP and demonstrated their viability and functionality.

## Results

### Dataset and Training

Two fine-tuning datasets were constructed: ft35k (35,150 plasmids, ≤100 kbp) and ft15k (14,485 plasmids, ≤30 kbp, circular only). Both datasets were restricted to *E. coli* plasmids, curated from PlasmidScope for natural plasmids and Addgene for synthetic (Table 2 (Methods)). Plasmid length distributions were highly skewed, with most plasmids below 10 kbp and a long tail towards the respective dataset cutoffs. Median plasmid length lies just above 5 kbp in both datasets (5681, 6488 bp), with wide interquartile ranges and a long tail towards the upper cutoffs. GC content distributions were tightly centered around ∼50%, consistent with the genomic composition of *E. coli* [28]. A small secondary spike around 40% GC was observed in the broader 35k dataset but diminished in the more restricted 15k dataset, suggesting that filtering out longer plasmids disproportionately removed low-GC constructs (Figure 1C). Topology filtering demonstrated that the 35k dataset contained a mixture of circular and incomplete/linear entries. Restricting to the 15k dataset reduced the number of circular plasmids by around 5’000, while eliminating the remaining linear/incomplete entries and thus represents the most biologically clean subset (Figure 1D).

**Figure 1:**
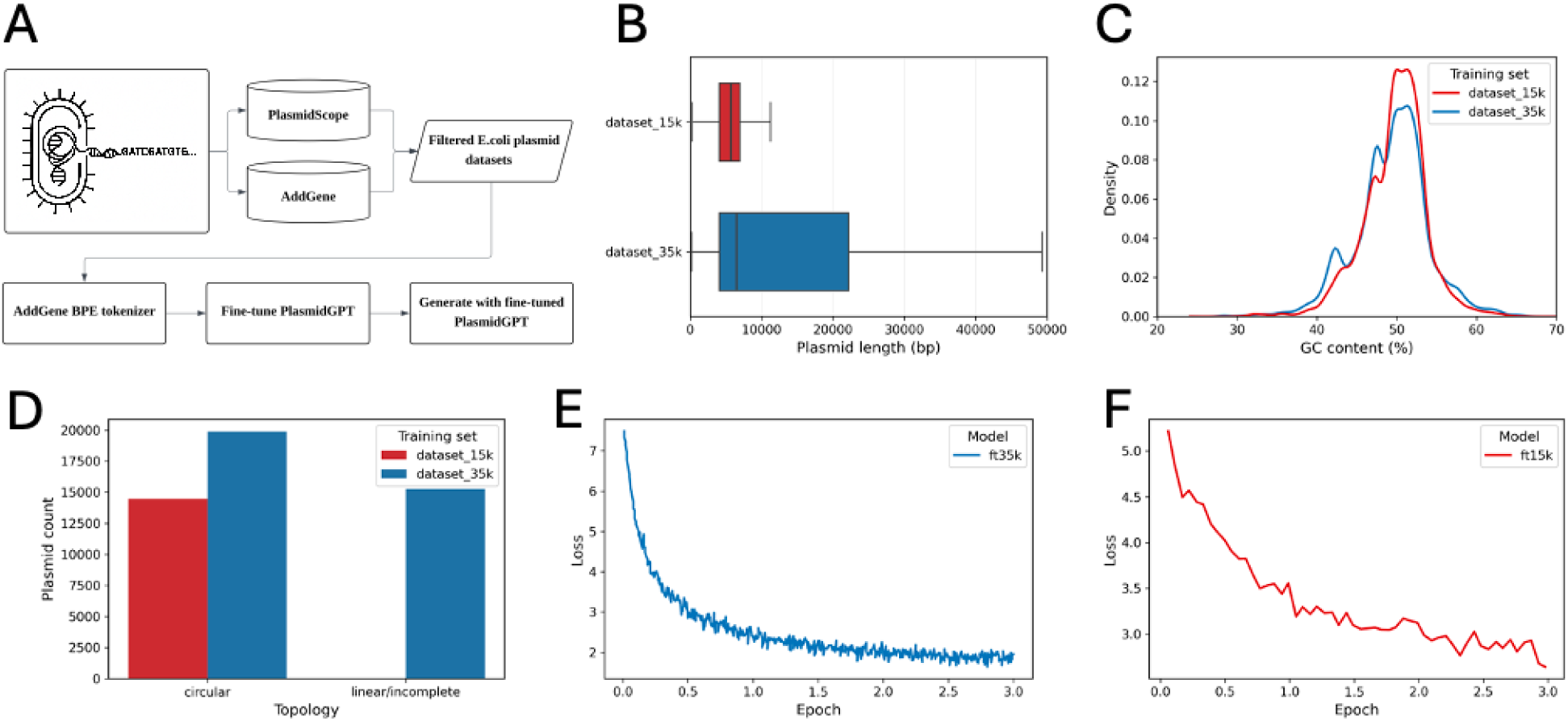
Curated *E. coli* plasmid data and fine tuning of models. A) End-to-end pipeline from data set curation from databases through to Addgene BPE tokenization, fine-tuning and generation with PlasmidGPT. B) Plasmid length distributions across the two fine-tuning datasets (dataset 15k, dataset 35k with medians of 5681, 6488 bp respectively). C) GC content (%) distributions across the two fine-tuning datasets. Addgene plasmids were included in each dataset. D) Counts of circular versus linear/incomplete plasmids across the two fine-tuning datasets. Addgene plasmids were assumed to be circular, while topologies from other databases were taken directly from metadata. E,F) Training loss curves for PlasmidGPT fine tuning on the 35k dataset (E) and the 15k dataset (F).

All fine-tuning experiments were conducted using PlasmidGPT, a GPT-2–style decoder-only language model pretrained on plasmid sequences with a 29,999-token BPE vocabulary. Fine-tuning was applied separately on the 35k and 15k datasets. Training curve for the 35k run showed smooth declines in loss across epochs, with no evidence of instability or divergence (Figure 1E). The ft15k run also converged successfully, dropping from ∼5.2 to ∼2.8 over three epochs, but the trajectory was somewhat less smooth (Figure 1F). This may be due to the reduced dataset size (14,485 vs. 27–35k plasmids), which limits the diversity of training batches and introduces more noise in gradient updates. Additionally, the random circular cropping used in the ft15k collator injects more stochasticity into the training signal than the deterministic sliding-window approach used in the 35k run. Overall, models trained stably, but the ft15k model benefitted from stricter biological filtering while showing slightly noisier optimisation dynamics.

### Analysis of Generated Plasmids

A total of 1,000 plasmids were generated from each model with a weak ATG seed and a stronger GFP expression cassette (Figure 2A). Generated lengths followed the triangular distribution used for targeting, centred on 4 kb with a spread of ±1.5 kb. The resulting density plots (Figure 2B) show consistent length distributions across all six sample sets (base ATG, base GFP, ft15 ATG, ft15 GFP, ft35 ATG, ft35 GFP), with the majority of plasmids clustering between 2.5 and 6 kb.

**Figure 2:**
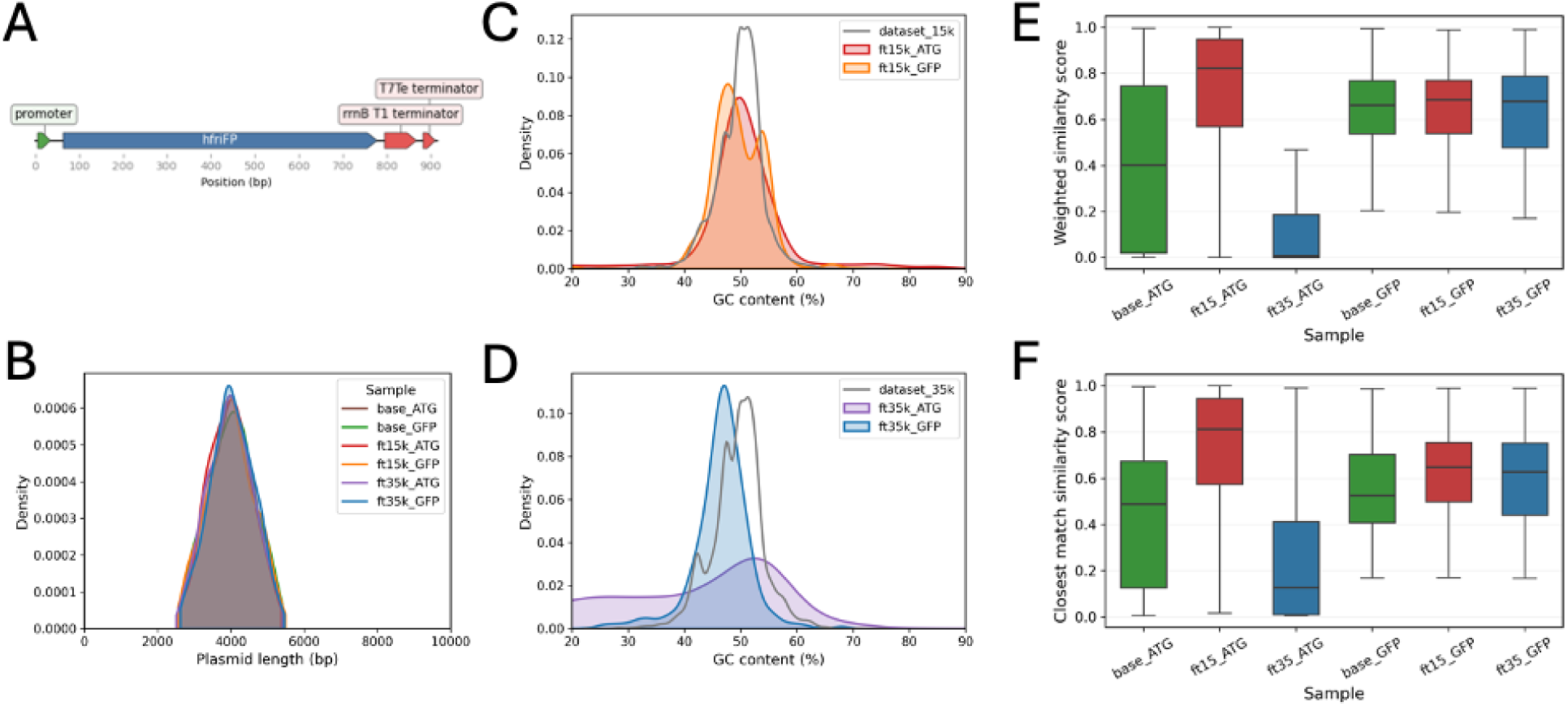
Analysis of generated plasmids. A) Annotated map of the GFP expression cassette sequence used for prompting strategy (917 bp). B) Density plots of generated plasmid lengths across six samples (base ATG, base GFP, ft15 ATG, ft15 GFP, ft35 ATG, ft35 GFP). C) GC content (%) for ft15k model generations (ATG and GFP seeds) compared with the ft15k training set. D) GC content (%) for ft35k model generations (ATG and GFP seeds) compared with the ft35k training set. E,F) Weighted true similarity scores and weighted closest-matched similarity scores across all six sample sets. Values represent aggregated BLAST identities and coverage across the full plasmid length relative to the training database.

Across all models and prompts, GC content distributions centred near 50%, consistent with the composition of *E. coli* (Figure 2C,D). Under ATG prompting, both the base and ft35k models exhibited broader GC spreads, with tails extending outside the desirable *E. coli* range. When GFP was used as the seed, these distributions tightened and clustered more closely around 50%. This contraction is expected: the GFP cassette imposes a coding context and local k-mer prior that anchors early generation toward typical *E. coli* codon usage, thereby reducing drift in GC during continuation. In contrast, the ft15k model maintained a consistently narrow GC spread (approximately 40–60%) under both ATG and GFP prompting, indicating that it internalised a stronger *E. coli* GC default during fine-tuning. Likely contributors include the more constrained 15k training set (restricted to circular-only, shorter backbones) and the training collator’s random circular crop, which uniformly exposed the model to composition across the plasmid and limited sensitivity to arbitrary start points. By comparison, ft35k was trained on a broader length distribution, plausibly inheriting more GC heterogeneity, which resurfaced under weak prompting (ATG) but was damped by the GFP seed.

With ATG prompting, both fine-tuned models exhibited lower JS and KL divergence to their respective training sets than the base model (Supplementary Figure S1) Reduced divergence indicates closer alignment to the empirical 4-mer distribution of plasmids, consistent with successful adaptation toward the target sequence manifold during fine-tuning. Using *k* = 4 captures context beyond single codons (*k* = 3), making the comparison more informative about local motif usage and short-range dependencies without incurring excessive sparsity.

Under ATG prompting, all models exhibited a broad spread of similarity scores, but with clear separation in central tendency (Figures 2E,F). The base model centered lower, with a median of approximately 0.4 for weighted true similarity and 0.5 for closest-match, indicating partial alignment to known backbones with sizeable unaligned regions. The ft35k model showed lower still (bulk around 0.2), consistent with either more aggressive novelty (mosaic assemblies that rarely resembled any single training plasmid over large spans) or drift off the training manifold under a weak seed. By contrast, the ft15k model scored high on both metrics (medians ∼0.8 for true and closest-match), implying strong, broad-coverage resemblance to training plasmids rather than isolated local matches; the small gap between closest-match and true similarity suggested that most of the sequence aligned well, not only a single region. With GFP prompting, similarity distributions converged across models toward a median of around 0.6 on both metrics, with tighter interquartile ranges. This homogenisation was expected: the shared cassette and associated promoter -RBS -terminator motifs increased alignable content and imposed similar local composition, thereby reducing between-model differences that were evident under ATG.

### Bioinformatics Assessment of Generated Plasmid Sequences

We next applied our bioinformatics pipeline to the generated plasmid sequences (Figure 3A). Under the ATG prompt, ORI counts were predominantly zero for the Base and ft35k models, whereas ft15k produced a markedly higher fraction of plasmids containing exactly one origin (Figure 3B). This likely reflects its circular-only, shorter-length training corpus and random circular cropping, which increased exposure to ORI contexts. In contrast, ft35k, trained on longer sequences with sliding windows, showed dilution of ORI motifs under weak prompting. Multiple-ORI outcomes were rare across all models, as expected for stable *E. coli* backbones. Introducing a GFP cassette shifted all models toward producing exactly one ORI (Figure 3C). The cassette provides a strong vector-like context that appears to cue the model to assemble a complete backbone, reducing reliance on model-specific priors. Although absolute ORI counts depend on detection thresholds, relative trends were maintained under stricter criteria.

**Figure 3:**
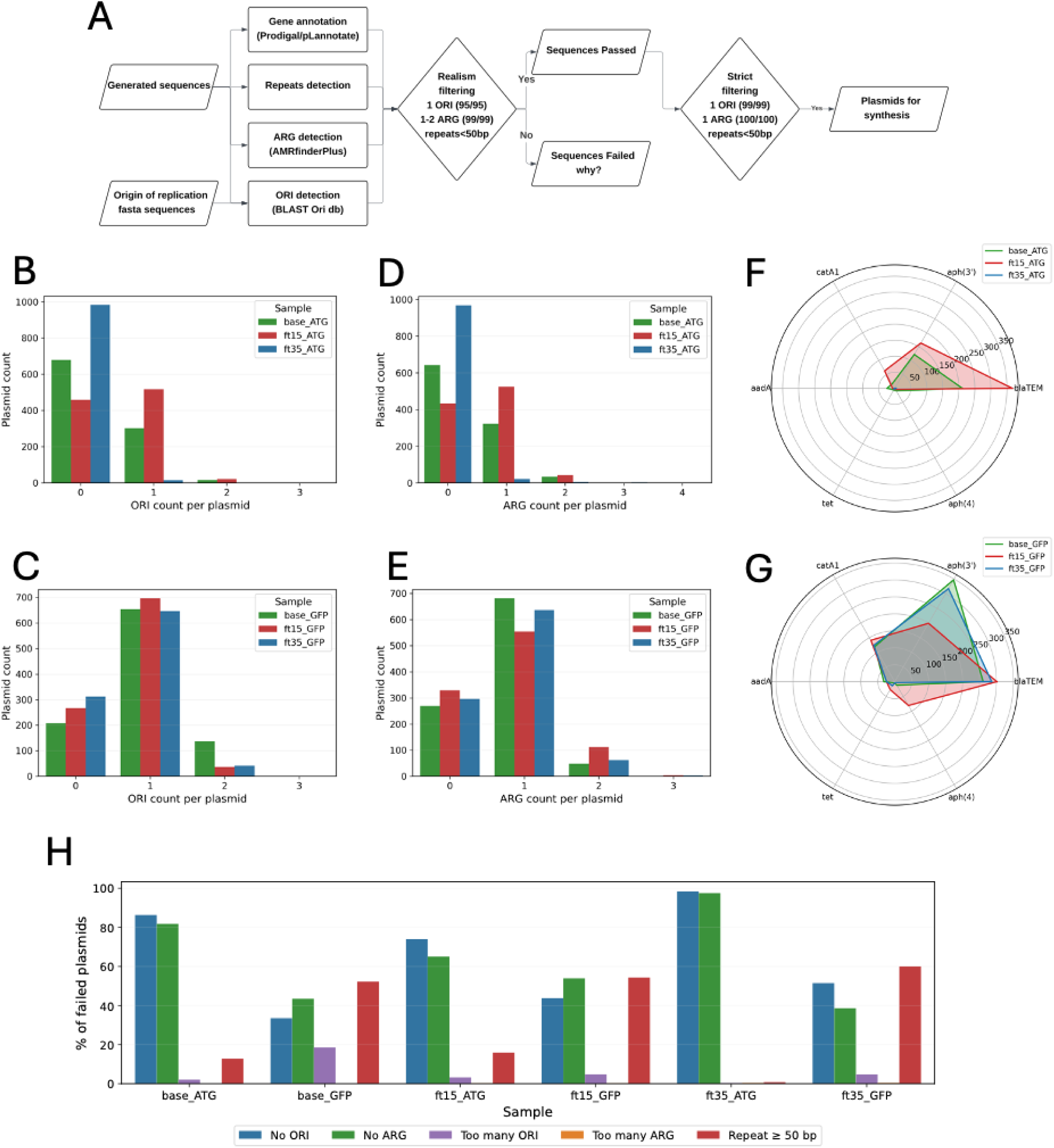
Bioinformatic assessment of generated plasmids. A) Bioinformatics pipeline from generated sequences to assessment (annotation, ORI/ARG typing, repeat detection); and two-tier screening (broad realism, then strict synthesis criteria) to select top backbones for wet-lab evaluation. B,C) Bar plots showing the distribution of origin of replication (ORI) counts in generated plasmids for ATG prompt and GFP prompt respectively. D,E) Bar plots showing the distribution of antimicrobial resistance (ARG) gene counts for the ATG prompt and GFP prompt respectively. F,G) Top six ARG families detected. Radar plots summarising the most frequent ARG classes detected across generated plasmids for the ATG-prompted and GFP prompted models respectively. Grouping was applied to collapse closely related variants (e.g. blaTEM-116 under blaTEM). H) Grouped bar plots of failure categories across all models and prompt types. Failures were grouped into six categories: No ORI, No ARG, Too many ORI, Too many ARG, and Repeat ≥50 bp. This highlights the dominant failure modes driving attrition during realism checks.

ARG counts paralleled ORI behaviour. Under ATG prompting, Base and ft35k rarely produced a selectable marker, while ft15k more consistently generated plasmids with exactly one ARG (Figure 3D). Instances of two ARGs were uncommon across models. With GFP prompting, all models shifted toward one ARG as the modal outcome (Figure 3E), again reflecting cassette-driven reinforcement of backbone structure. ARG composition further highlighted these trends (Figure 3F,G). Under ATG, Base and ft15k outputs were dominated by *blaTEM*, with secondary aminoglycoside and chloramphenicol markers, whereas ft35k produced sparser assignments. GFP prompting shifted Base and ft35k toward aph(3), consistent with common GFP+kan backbones [3]. ft15k retained the broadest repertoire, including occasional aph(4), reflecting its cleaner, biologically focused training set. Repeat distributions are shown in Figure S2. Most plasmids contained longest repeats ≥ 50 bp, below the instability threshold used in filtering. Under GFP prompting, a heavier right tail emerged, likely due to cassette-adjacent motif echoing and the mild decoding temperature. These longer repeats were removed in downstream filtering, affecting pre-filter distributions but not the final candidate pool.

### Filtering and Experimental Realisation

Table 1 summarises the proportion of generated plasmids satisfying the broad realism filter, defined as exactly one origin of replication (ORI) and one antimicrobial resistance gene (ARG), with high-confidence thresholds of ≥ 95% identity and coverage for ORIs and ≥ 99% identity and coverage for ARGs. These criteria reflect the requirement for intact, functional backbone components suitable for potential synthesis.

**Table 1:** Proportion of generated plasmids passing realism criteria. Summary of the percentage of plasmids (out of 1000 generated per model/prompt) that satisfied the basic realism filter: exactly one origin of replication (ORI) and one or two antimicrobial resistance (ARG) genes, both meeting identity and coverage thresholds. The table compares Base, ft15k, and ft35k models under both ATG and GFP prompts.

| Model | Prompt | Pass Rate (%) |
| --- | --- | --- |
| Base | ATG | 17.9 |
| Base | GFP | 23.9 |
| ft15k | ATG | 27.6 |
| ft15k | GFP | 19.0 |
| ft35k | ATG | 0.1 |
| ft35k | GFP | 11.1 |

**Table 2:** Number of plasmids included in each fine-tuning dataset by source. Sources with zero contribution across all datasets were omitted for clarity.

| Dataset | GenBank | RefSeq | PLSDB | IMGPR | DDBJ | COMPASS | Addgene | Total |
| --- | --- | --- | --- | --- | --- | --- | --- | --- |
| ft35k | 14,878 | 1,892 | 624 | 5,047 | 1,245 | 1,160 | 10,304 | **35,150** |
| ft15k | 3,022 | 245 | 211 | 1 | 368 | 334 | 10,304 | **14,485** |

Under the minimal ATG seed, pass rates diverge sharply across models. The ft15k model achieves the highest rate at 27.6%, followed by the base model at 17.9%, while ft35k performs near zero at only 0.1%. This ordering reflects differences in training corpora. ft15k, fine-tuned on a circular-only, shorter-length dataset with random circular crops, more consistently inserts both an ORI and a selectable marker that meet the strict thresholds. The base model more often omits one of these features or produces a sub-threshold variant, reducing its pass rate. ft35k, trained on longer plasmids, dilutes ORI and ARG motifs across wider contexts, explaining the near-total absence of valid backbones under weak prompting.

When a GFP cassette is used as the seed, the ordering shifts. The base model rises to a 23.9% pass rate, outperforming ft15k at 19.0%, while ft35k improves substantially to 11.1%. The GFP cassette acts as a strong vector-like prior, steering all models toward canonical repository patterns such as GFP+kan or GFP+amp backbones. Under this conditioning, the broadly pretrained base model snaps more frequently to clean 1-ORI / 1-ARG designs above threshold. ft15k remains disciplined but is occasionally penalised at the margins: either through ARG variants falling just below the 99/99 cutoff, or cassette-adjacent duplications that increase repeat burden if included in the realism screen. As a result, the advantage it held under ATG is diminished under GFP, with the base model edging ahead. ft35k benefits most from GFP conditioning, though its backbone prior remains weaker overall.

Failure-mode analysis provides further insight into these patterns (Figure 3H). Under ATG prompting, the dominant reasons for attrition are missing ORIs and ARGs, consistent with weak seeding that fails to cue complete backbone assembly. By contrast, under GFP seeding, the principal failure category shifts to excessive repeats, a side-effect of cassette-conditioned motif echoing and occasional local duplication. Comparing models, the base and ft15k show broadly similar distributions of failure categories. The main distinction is that the base model produces too many ORIs in roughly 15% of cases, whereas ft15k rarely makes this error, reflecting its stricter circular-only training prior. Overall, repeat sequences emerge as the chief bottleneck for GFP-conditioned designs, while incomplete backbones dominate the ATG regime.

### Synthesis Candidates

The ft15k model was selected as the basis for candidate backbones because it was fine-tuned on a circular-only corpus restricted to plasmids ≤ 30 kb, strengthening its backbone priors and limiting drift. Compared with the base and ft35k models, ft15k generations exhibited tighter GC distributions and higher realism under weak (ATG) prompting. Its use of random circular crops during training also increased exposure to essential motifs such as ORIs and ARGs across all sequence contexts, while the shorter, cleaner completions it produced aligned well with synthesis constraints.

Generations were then subjected to the synthesis-grade filtering and prioritisation workflow (Methods). To survive this stage, each plasmid was required to contain exactly one ORI at ≥ 99% identity and coverage, exactly one ARG at 100% identity and coverage, and no internal repeats ≥ 50 bp. The pool was further constrained to plasmids shorter than 4,000 bp to reduce synthesis cost, and restricted to high-copy, self-replicating backbones (e.g. ColE1-family derivatives) to avoid helper requirements. This pipeline yielded 16 plasmids in total. Candidates were ranked by minimal repeat burden (shortest longest repeat), balanced novelty (reduced weighted similarity when compared against full NCBI database), and lean annotation profiles (few or no extraneous features beyond ORI, ARG, and GFP). The top three were selected for further analysis.

The four final candidates each contain a chloramphenicol-resistance gene (*catA*) and a ColE1-like origin of replication, but differ in the spatial arrangement of these features relative to the GFP cassette. Intervening sequence segments exhibit sparse minimal annotations under pLannotate, suggesting compact non-coding scaffolds and/or regulatory DNA not well represented in reference libraries. While unannotated, such regions may influence copy number, transcriptional interference, and stability. They were retained on the basis that they passed all strict criteria, exhibited low repeat burden, and maintained balanced novelty. Representative maps of the candidates are shown in Figure 4. An example of a failed plasmid is is displayed in Supplementary Figure S3 which contained a window of low GC content (see 1600-1900bp region) and was not taken forward for synthesis and construction.

**Figure 4:**
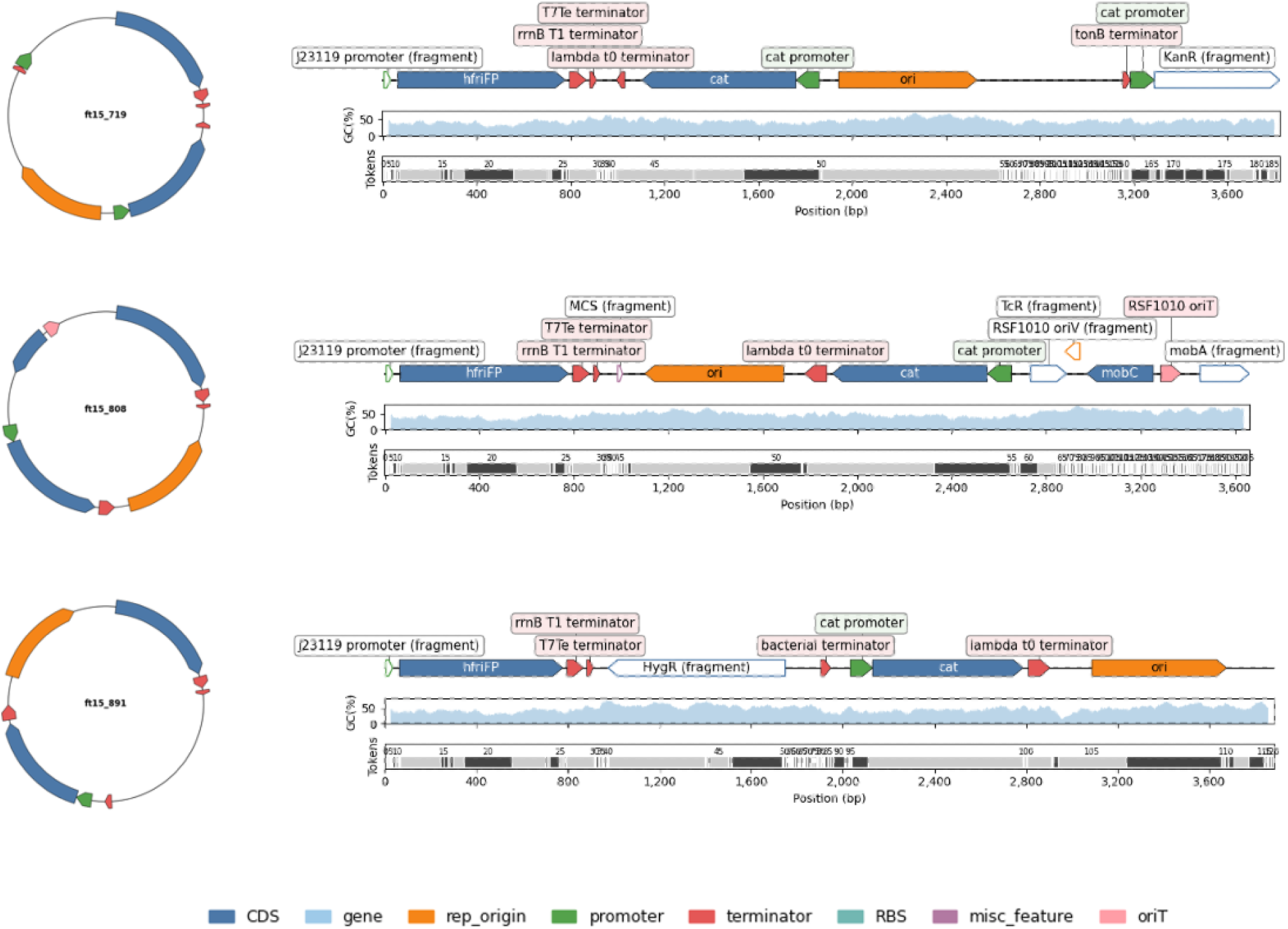
Realisation of plasmid designs. Representative circular and linear backbone maps for the synthesis candidates (ft15 719, ft15 808, ft15 891). These plasmids satisfy the criteria of exactly one ORI, one ARG, and no repeats ≥ 50 bp. GC content(%) and token alignment displayed for each candidate.

### Construction and Characterisation of Generated Plasmids

The three remaining candidates, 719, 808, 891 were synthesised and constructed using standard assembly approaches and then characterised. These constructs all successfully grew, produced functional antibiotic resistance and expressed GFP (Figure 5). Most strains grew to similar OD levels but with different GFP levels compared to the positive control formed from the same GFP casette in the standard MoClo backbone. All colonies produced at least as much GFP expression as the control if not more. These results confirm that all three AI-generated plasmid backbones were not only viable but fully functional in vivo, demonstrating that the model can produce operational genetic constructs.

**Figure 5:**
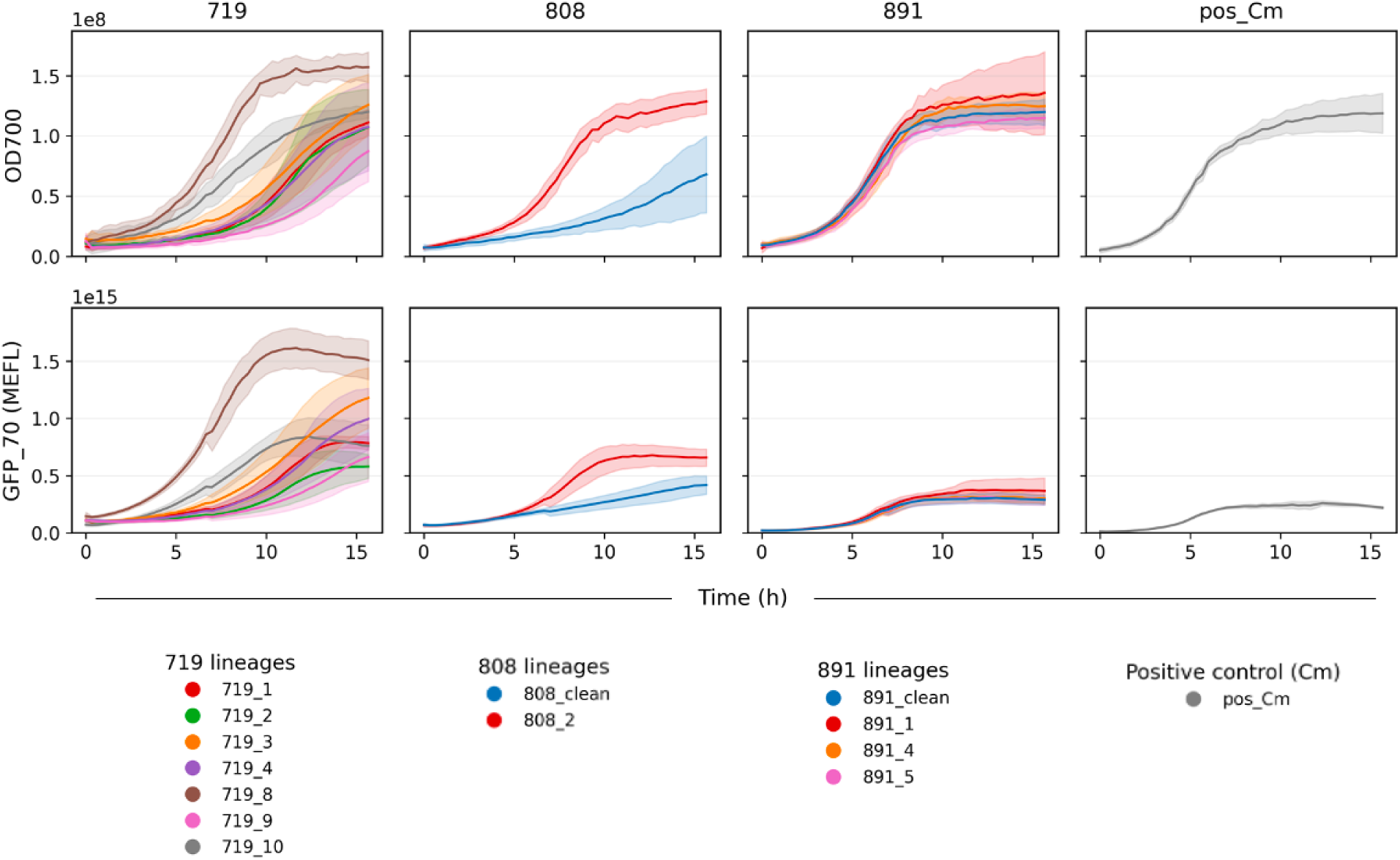
Growth dynamics and GFP fluorescence across engineered plasmid variants. Time-course measurements of optical density representity relative cell count (OD700, top panels) and GFP expression in MEFL (bottom panels) for colonies harbouring different plasmid backbones. Each column represents a distinct construct group (ft15 719, ft15 808, ft15 891, and a positive control), with individual mutated colonies shown as separate traces ( clean representating no mutations). Positive control (Standard MoClo Backbone) samples are included to benchmark expression and background signal.

Interestingly, there were stark differences in mutational stability across the plasmids (Figure S3). Plasmid 719 had mutated in 7/7 colonies sequenced, which could be due to high copy-number/inverse repeat of cat promoter. Plasmid 808 had mutated in 1/5 colonies sequenced. Plasmid 891 had mutated in 3/5 colonies. Most mutations were small indels or point substitutions localised near regulatory regions rather than across the GFP cassette. This mutational heterogeneity caused varied performance in terms of OD and GFP expression, in some cases causing higher OD and GFP expression than the control (e.g. 719 8, 808 2, 891 1). The emergence of distinct mutational profiles across otherwise similar backbones highlights that generatively designed plasmids can be functional yet evolutionarily dynamic. This reflects genuine biological pressures acting on replication, burden, and regulatory architecture which reveals sequence features that future models may need to account for explicitly.

## Discussion

This study is the first time a DNA large language model has been used to design and construct new plasmids that are functional. This was achieved by establishing a full pipeline for plasmid backbone generation and assessment using PlasmidGPT, from dataset curation through strict candidate selection. Two *E. coli* datasets were prepared (35k and 15k), with the 15k set (14,485 circular plasmids ≤ 30 kb) providing the cleanest training corpus. Fine-tuning employed circular-aware batching and random crops; all models converged, though ft15k exhibited noisier loss trajectories consistent with its smaller size. Generations (1,000 per model/prompt) were produced under two seeds: a weak ATG and a functional GFP cassette. Lengths and GC distributions generally matched *E. coli* norms; ft15k maintained the tightest GC envelope across prompts. Fine-tuned models reduced 4-mer divergence relative to Base, and similarity analyses showed clear separation under ATG (ft15k highest, Base intermediate, ft35k lowest), with convergence around 0.6 under GFP. Backbone assessments highlighted that ft15k more reliably produced single-ORI/single-ARG plasmids under ATG, while GFP seeding regularised all models toward canonical backbones. ARG composition reflected common markers (*blaTEM*, *aph*(3^′^)), with repeats generally *<*50 bp but longer tails emerging under GFP. Broad realism filtering (one ORI, one/two ARGs, repeat *<*50 bp) yielded pass rates of 18-28% for Base/ft15k and ≤12% for ft35k, with ft15k outperforming under ATG and Base performing best under GFP. Stricter synthesis-grade filtering reduced candidates to 16, from which four ft15 GFP plasmids (ColE1 + *catA*) were shortlisted for wet-lab testing, three of which made it to synthesis and construction. The three AI generated plasmids grew, produced antibiotic resistance and expressed GFP when compared to the control. However, they did show mutational instability with mutations found in the generated origin of replication, antibiotic resistance gene and in the backbone.

Beyond these quantitative assessments, the experimental data revealed several biological behaviours that provide insight into both the strengths and current limitations of generative plasmid design. The observed mutational instability is interesting from a biological point of view. Some mutations are in the origin, which are likely to alter the copy number of the plasmids. They all express the same cassette and have highly similar origins (*>* 99% similarity), but the biology of plasmid replication is complex and could explain these differences [23]. It is noted that mutations were also common in the synthesis of AI designed phages [21]. This all points to the fact that there is the potential for optimising plasmid backbones using our approach, which we will address in future work. There are a number of other limitations, which can be improved in future version. The Addgene-trained BPE tokenizer (29,999 tokens) contains a long tail of unusually large subwords (696 tokens *>*400 bp), including entire functional parts with 131 tokens containing an ORI and 169 tokens containing an ARG (85% identity and 80% coverage). Such “macro-tokens” allow the model to place large, high-level modules in a single step, which blurs local base-pair dependencies, artificially inflates realism pass rates, confounds novelty analysis, and risks evaluation leakage if token content overlaps with training or reference databases. This could be remedied in a number of ways: either capping the maximum token length (e.g., 3-6 bp); retraining the BPE/unigram to prevent macro-tokens; and exploring hybrid schemes (character-level for background, short subwords for motifs, or k-mer tokenisation with circular rotation). Assessment of plasmids relied heavily on BLAST, AMRFinderPlus, and pLannotate. These alignment-based methods perform well for known parts but miss weakly conserved motifs, novel ORFs, and unannotated regulatory signals in spacer regions. Weighted similarity aggregates alignments but underweights unaligned, potentially novel content. In terms of prompts, GFP seeding increased the right tail of repeat lengths (≥50 bp), likely reflecting motif echoing around promoter/terminator/UTR segments under stochastic decoding. Although downstream filtering removed these cases, the effect reduced yield and biased selection toward certain layouts. Finally, although a large training set was established, access to more and greater diversity of sequences would further enhance the novelty of the generations and allow for a complete training of a more general Large Language model. Future work will also include the extension to other hosts with tagged corpora, and add lightweight control tokens (e.g., <ONE ORI>, <ONE ARG>, <HIGH COPY>) to test controllability without fully specifying seeds and the use of larger models such as Evo 2. The future will see the use of reinforcement to further fine tune models. This can be done bioinformatically, but closing the loop by incorporating wet-lab results into model refinement would be advantageous. Together, these directions lay the foundation for a more automated plasmid engineering workflow, where generative models incorporate host context, design constraints, and experimental fitness signals to refine backbone architecture over successive iterations. Such integration of controllable generation, multi host training, and reinforcement from wet-lab measurements would enable plasmids to be optimised not only for functionality but also for stability and manufacturability.

In summary, we have demonstrated for the first time how DNA language models can be used to generate viable sequences and experimentally testable plasmid designs, through data curation, fine-tuning, and prompt-aware generation. Our approach opens up the possibility of using AI in the design of DNA sequences for broader applications across engineering biology.

## Methods

### Dataset Curation

#### Sources and Scope

Plasmid sequences were obtained from PlasmidScope as of June 2025 [24], which aggregates across multiple databases including GenBank [29], RefSeq [30], PLSDB [31], IMG-PR [32], DDBJ [33], COMPASS [34] and Kraken2 [35]. This yielded approximately 852,600 plasmid records. To maintain host-specific relevance, only plasmids annotated for *Escherichia coli* were retained, resulting in a final collection of 29,608 sequences. In parallel, 10,304 plasmids were incorporated from the Addgene repository [3], previously scraped for bacterial expression backbones (up to september 2024). This created a full dataset of 39,912 sequences. Where overlaps occurred across databases, duplicates were resolved by keeping the first identifier alphabetically by database. This ensured consistent deduplication without inflating plasmid counts. Table 2 summarises the contributions of each source to the final fine-tuning datasets.

#### Preprocessing and Filtering Strategy

To construct datasets of manageable size and biological plausibility, progressive filtering was applied. Both datasets included the full previously scraped Addgene plasmids.

**ft35k (35,150 plasmids):** Retained all *E. coli* plasmids ≤100 kbp. This was done in order to include as much genetic data as possible without the interference of large sequences dominating the training. This set provided the broadest coverage but included long-tailed length distributions.

**ft15k (14,485 plasmids):** Subset of the above, restricted to plasmids ≤30 kbp. This removed larger complexes dominating the training. Additionally, only plasmids explicitly annotated as circular were retained. Linear or incomplete entries were removed, along with entries flagged as “fragments” or potential phages. This dataset formed the final training set, optimised for a goal of full plasmid design (full circular plasmids only) and computational tractability (shorter maximum length).

### Model Training

#### Tokenisation and Circular Handling

Tokenisation was performed using the byte-pair encoding (BPE) tokenizer created with PlasmidGPT, trained on large-scale plasmid corpora and containing a vocabulary of 29,999 tokens. This ensured compatibility with the pretrained model. Unlike natural language tasks, no beginning-of-sequence (BOS) or end-of-sequence (EOS) tokens were introduced. This was to reflect the circular nature of plasmids, which lack natural start and end points. Forcing an EOS token would have created artificial termination signals with no biological counterpart.

To handle circularity, a circular cropping strategy was applied. Each training example consisted of a 2,048-token segment randomly cropped from the plasmid sequence, with sequences duplicated *in silico* to allow wrap-around. This ensured that the model was exposed to all possible subsequences without creating an arbitrary cut-point. For the 35k dataset, tokenisation used a sliding-window approach with stride 1,024, producing overlapping 2,048-token contexts. For the 15k dataset, tokenisation was applied to full plasmid sequences, with a single random 2,048-token circular crop taken per plasmid per step. Shorter plasmids (*<*2,048 tokens) were included in full and dynamically padded within batches.

To reduce unnecessary computation on padding tokens, batches were bucketed by length so that plasmids of similar size were grouped together before padding. For the 15k dataset in particular, padded tokens were explicitly ignored in the loss function: attention masks set padded positions to –100 in the labels, ensuring they did not contribute to gradient updates. This collator design allowed efficient training on highly variable-length plasmids while preserving correct handling of circular crops.

#### Base Model

The foundation for all fine-tuning was PlasmidGPT, a decoder-only, GPT-2-style language model specifically trained to model plasmid DNA [22]. PlasmidGPT was introduced in 2024 as a model capable of learning the “sequence grammar” of plasmids, including origin of replication (ORI) types, backbone functions, and motifs commonly found in natural plasmids. Key features of the model include pretraining on a vast set of plasmids from multiple sources, giving exposure to diverse ORIs, promoters, repeats, and other elements, and the use of BPE tokenisation with a large vocabulary that allows efficient representation of frequent subsequences such as codon combinations or regulatory motifs. For this project, the pretrained PlasmidGPT model and its associated tokenizer (vocabulary size = 29,999 tokens) were reused directly. This enabled fine-tuning to build on learned representations rather than training from scratch.

#### Fine-Tuning

Fine-tuning was applied to the datasets (35k and 15k) to assess how sequence length constraints and topology filtering affect PlasmidGPTs adaptation for generation. Training was performed with a context length of 2,048 tokens, a batch size of one sequence per device with gradient accumulation over eight steps, a learning rate of 5 × 10^−5^, and a total of three epochs. Five hundred warmup steps were used, and checkpoints were saved every 500 steps, retaining the two most recent per run. For the 35k dataset, sliding-window tokenisation with stride 1,024 was used, producing overlapping contexts. For the 15k dataset, tokenisation was applied to full sequences and, at each batch, a random circular crop was taken, or the entire sequence was retained with padding if shorter. Padded positions were masked out of the loss function as described above. Fine-tunes were run on GPUs (RTX 3070 and HPC nodes with equivalent resources). Gradient checkpointing was enabled where required to balance compute and memory usage, allowing efficient training across all dataset scales.

### Sampling Procedure (Generation Focus)

#### Prompting Strategy

For each model variant (base, ft35k, ft15k), a total of 1,000 sequences were generated. Two seeding strategies were employed to initiate sampling. The first used a curated GFP expression cassette as a conditioning scaffold displayed in Figure 2A. This cassette provided promoter-RBS-GFP-terminator motifs and served three purposes: it biased generation toward GFP-ready plasmid backbones consistent with the project aim, it offered a biologically meaningful context around which downstream analyses could assess coherence of origins, ARGs, and backbone structures, and it enabled controlled comparisons against unprompted sampling. The second strategy used a minimal seed consisting only of the trinucleotide ATG. This prompt was selected because it corresponds to the canonical start codon in bacteria, providing a weak but biologically meaningful initiation signal [36]. This avoids biasing the model with vector-like context (as in GFP prompting) while still anchoring generation in a plausible coding motif. Alternatives such as ATGC were avoided, as they introduce arbitrary bias (a codon plus partial context) without clear biological justification and may skew local *k*-mer composition in unintended ways. All sampling employed the PlasmidGPT BPE tokenizer to ensure compatibility with fine-tuned checkpoints.

#### Length Targeting and Stopping Criteria

A triangular distribution was chosen to target plasmid lengths because it provides bounded support and explicit control over the mode. This ensured sampled targets remained within practical synthesis limits (2500–5500 bp), while biasing toward the common 4 kb backbone size. In contrast, a normal distribution has unbounded tails and would frequently produce implausibly short or long plasmids unless truncated, wasting samples or overweighting extremes. The distribution was parameterised such that

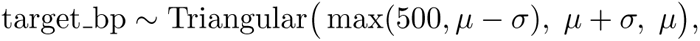

where *µ* = 4000 and *σ* = 1500. Generation was halted according to a base-pair budget criterion implemented as a custom stopping criterion. For each token, the decoded bases were counted, and the cumulative number of emitted nucleotides was tracked relative to the prompt. Once this total exceeded the sampled target length, generation was stopped. A conservative ceiling of 12,000 new tokens was also applied, but this limit was rarely approached due to the bp-budget stopping rule.

#### Decoding Configuration

Decoding parameters were selected based on preliminary sweeps that balanced sequence diversity against biological plausibility. All final runs used a temperature of 1.05, nucleus sampling with top-*p* = 0.95, and a repetition penalty of 1.05. No repeat-*n*-gram constraint was applied, this was because tokens varied widely in bp length and so there was no obvious constraint to give. No EOS token was supplied, since plasmids are circular. These settings were held constant across all models to ensure comparability.

#### Post-processing and Circular Trimming

Raw decoded sequences were sanitised to the IUPAC DNA alphabet [ACGTN]. To prevent overshoot and approximate circular closure, a post-processing function was applied. This function anchored on the first 64 bp of the generated sequence and searched for its next occurrence downstream, requiring that the match appear beyond a minimum of 60% of the target length but within 140% of it. If such a return point was identified, the sequence was trimmed at this position to emulate circular wrap-around. If no repeat was found, the sequence was truncated to remain within plausible plasmid sizes, with enforced bounds between 1,000 bp and the target length. A binary flag recorded whether circular trimming was applied, ensuring traceability of post-processing outcomes.

### Bioinformatics Assessment Pipeline

The generated plasmids were systematically evaluated using a multi-stage bioinformatics pipeline designed to assess realism and novelty. The pipeline combined gene annotation, replicon typing, antimicrobial resistance (ARG) detection, repeat screening, and similarity analysis.

#### Annotation and Gene Prediction

Prodigal was used for rapid gene calling to identify potential coding sequences, including novel constructs not captured by curated references. This provided a lightweight first-pass annotation of protein-coding regions. In parallel, pLannotate was employed for cross-referencing and visualisation. pLannotate integrates multiple sequence databases, including those for resistance genes, replication origins, and functional domains, and produces annotated plasmid maps that facilitated manual inspection of generated backbones.

#### Origin of Replication Detection and Typing

Origins of replication were identified by BLAST alignment against a curated ORI reference database compiled from NCBI entries and augmented with commonly used replicons in design such as ColE1. For each hit, both identity (%) and coverage (%) were recorded, and in cases of overlapping alignments the match with the highest identity was retained. Metrics collected for each plasmid included the ORI type, identity and coverage scores, genomic position, and the count of unique ORIs detected.

#### Antimicrobial Resistance Gene Detection

ARG content was assessed using AMRFinderPlus, which applies BLAST-based detection against curated resistance gene databases. As with ORI detection, identity and coverage were recorded for each hit, and the best alignment was retained where overlaps occurred. Metrics for each plasmid included ARG type or classification, identity and coverage scores, genomic positions, and the total number of ARGs identified.

#### Repeat Detection

To identify internal repeats that might destabilise plasmids or promote recombination, each sequence was aligned against itself using BLAST, with circularity taken into account. Repeats were extracted and scored by length, and the longest repeat region per plasmid was retained as a summary metric.

#### Divergence Metrics

To quantify how closely generated plasmids resembled training distributions at the compositional level, the Kullback–Leibler (KL) divergence and Jensen–Shannon (JS) divergence was computed between empirical *k*-mer frequency profiles [37, 38]. Let *P* and *Q* denote two discrete probability distributions over the same support (here, the set of all *k*-mers). The KL divergence of *Q* from *P* is defined as

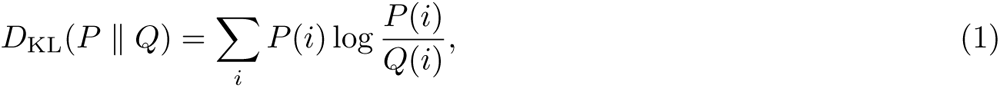

where *P*(*i*) and *Q*(*i*) denote the probability of observing the *i*-th *k*-mer under *P* and *Q*, respectively. KL divergence is asymmetric and non-negative, with *D*_KL_(*P* ∥ *Q*) = 0 if and only if *P* = *Q*. Because KL can diverge when *Q*(*i*) = 0, we also report the JS divergence, a symmetric and smoothed variant defined as

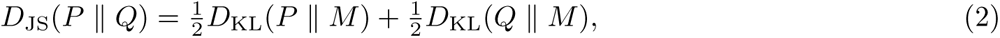

Where *N*is the mean distribution. JS divergence is bounded between 0 and 1 when base-2 logarithms are used, and equals 0 only when *P* = *Q*. In practice, we estimated *P* and *Q* as normalised frequency distributions of 4-mers from the training corpus and generated plasmid sets. Lower divergence values indicate closer alignment to training statistics, while higher values reflect compositional drift.

#### Novelty Assessment

Two complementary similarity analyses were implemented to evaluate novelty relative to the training set. First, direct BLAST searches against the training plasmids identified the best-matching sequence for each generated plasmid, allowing classification as a near-duplicate, a variant, or a novel construct. Second, a weighted similarity score was calculated to account for the modular nature of plasmids, where different regions may align to different training sequences:

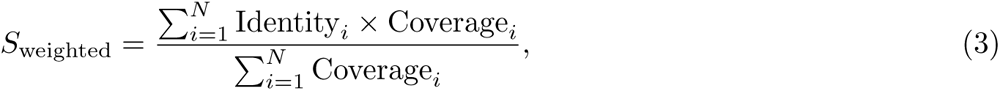

where *N* is the number of BLAST matches, Identity*_i_* is the percentage identity of the *i*th hit, and Coverage*_i_* is the proportion of the plasmid covered by that hit. This formulation weights identity by coverage, yielding a single score that reflects both sequence similarity and breadth of alignment. A high score dominated by a single source suggested a near-copy, whereas moderate scores distributed across multiple sources indicated recombination of modules from different plasmids. In addition, a best-match analysis recorded the source sequence of the top hit, the maximum local identity observed, and the training plasmid identifier corresponding to the strongest alignment. Together, these analyses revealed whether a generated plasmid represented a direct reproduction of a known sequence, a mosaic assembly of existing regions, or a genuinely novel combination.

#### Thresholds and Parameters

For all BLAST- and ARG-related analyses, thresholds were set at ≥ 85% identity and ≥ 80% coverage. These looser cutoffs were chosen to tolerate database variance and novel creations while maintaining meaningful similarity constraints. Where overlapping hits were identified, the alignment with the highest identity was retained. The full set of metrics collected for each plasmid included ORI types, identities, coverages, and positions; ARG types, identities, coverages, and positions; ORI and ARG counts; predicted coding sequences from Prodigal; longest repeated region; and the weighted similarity score and best-match identifier relative to the training data.

### Filtering and Candidate Selection

#### Realism Filtering

To identify plasmids that were biologically plausible and functionally viable, an initial round of broad filtering was applied. These thresholds were designed to reflect established principles in engineering biology while tolerating natural variation. Each plasmid was required to contain exactly one origin of replication (ORI), detected at a threshold of ≥ 95% identity and ≥ 95% coverage. Multiple replication origins are well known to destabilise plasmids and complicate copy number control, whereas allowing moderate sequence variability accommodates the natural diversity observed among replicons such as ColE1 and pMB1 [39]. Antimicrobial resistance genes (ARGs) were restricted to one or two per plasmid, with stricter detection thresholds of ≥ 99% identity and ≥ 99% coverage. These criteria reflect the functional constraints of ARGs, as even minor single point mutations can corrupt the genes, making near-perfect matches essential for biological relevance. In addition, plasmids containing internal repeats longer than 50 bp were excluded, since repeats of this magnitude are sufficient to act as substrates for homologous recombination, leading to structural instability, deletion, or rearrangement [40, 41]. The 50 bp cutoff was selected as a conservative threshold commonly cited in the literature on plasmid stability. Other constraints on plasmid length, GC content, and topology had already been enforced during dataset curation and generation, *E. coli*-compatible plasmids were assessed. This broad filtering stage produced a pool of sequences that were viable in principle: each contained a single replicon, intact ARGs, and limited recombinational risk.

#### Selection of ideal GFP backbone candidates for wetlab evaluation

From this pool, a second, more stringent filtering stage was applied to identify a small number of top candidates suitable for experimental synthesis. This case study imposed tighter thresholds to minimise ambiguity and maximise reliability. Exactly one ORI was required per plasmid, detected at ≥ 99% identity and ≥ 99% coverage, ensuring functional replication origins with virtually no divergence from reference sequences. For ARGs, the requirement was even stricter: exactly one ARG detected at 100% identity and 100% coverage. This absolute constraint eliminated the risk of truncated or inactive variants. Candidate plasmids were then ranked by lowest internal repeat burden, as reduced duplication decreases recombinational instability and facilitates reliable synthesis. To balance novelty against plausibility, plasmids were also ranked by weighted similarity score relative to the training dataset and against full NCBI database. This prioritisation avoided trivial reproductions of training sequences, while also excluding constructs so divergent that they might fall outside known biological plausibility. By combining these criteria, a ranking scheme was established that jointly optimised replicon integrity, ARG reliability, structural stability, and novelty. Plasmid maps were visually inspected on pLannnotate with a low threshold (coverage ≥ 10%) and SnapGene to check for plausibility and the top four plasmids were selected based as exemplars for downstream analysis and potential laboratory synthesis.

### Experimental methods

#### Plasmid construction

All generated plasmid backbone sequences were purchased as gBlock fragments (Twist Bioscience). The original GFP producing plasmid containing the J23106 constitutive promoter, B0034m RBS, E0040m (GFP) CDS and B0015 terminator, was constructed using DNA parts from the standard CIDAR MoClo kit [42] and inserted into a DVC261 AE backbone (kindly supplied by Casey Chen). The GFP cassette fragment was PCR amplified using Q5^®^ High-Fidelity DNA Polymerase and inserted into the synthesised plasmid backbone via NEBuilder^®^ HiFi DNA assembly (New England Biolabs) according to the manufacturers specifications. Cloning was performed in commercial NEB®5-*α* competent *E. coli* (New England Biolabs) and plasmid sequences confirmed with full-length plasmid sequencing (FullCircle). Correct plasmids were stored as glycerol stocks at −80^◦^C until needed.

#### Plasmid characterisation

Strains were inoculated from glycerol stocks and grown in M9 media for 18 h at 37 °C with shaking. Cultures were then adjusted to OD700 0.05 in fresh M9 media. 125 *µ*l of each diluted culture was added directly to the well of a 96-well plate, and the plate sealed with a Breathe-Easy^®^ sealing membrane. Plates were then incubated for 16 h at 37 °C with shaking (300 rpm, 2-mm-orbital), in a Tecan Spark plate reader (Tecan, USA). Measurements for OD700 and GFP fluorescence (excitation: 488/20 nm, emission: 530/20 nm, gain: 70) were collected every 20 min. The growth and fluorescence were then estimated using the FlopR package. A detailed description of this process is given by Fedorec et al. [43].

#### Strains and Plasmids

A full list of all reagents used in the manuscript are provided in Table S1. All strains and plasmids used in this study are given in Table S2. Primers used to create the HiFi fragments are given in Table S3.

### Biosafety Considerations

All work in this study was conducted under appropriate biosafety oversight and exclusively involved non-pathogenic *Escherichia coli* cloning strains and standard bacterial expression plasmids. Sequence data used for model training and generation were from public repositories and restricted to plasmids intended for laboratory cloning and protein expression, and did not contain pathogenicity islands, virulence factors, toxin genes, or sequences associated with regulated biological agents. No mammalian, viral, or eukaryotic pathogenic DNA was included. All experimental work employed commercially available *E. coli* NEB 5-*α* cells in containment level 1 (CL1) facilities under institutional biosafety procedures. The generative framework was therefore developed and evaluated entirely within a low-risk biological context, and all synthesised plasmids consisted solely of safe, well-characterised parts widely used in synthetic biology.

## Data and Code Availability

All code used for dataset construction, model fine-tuning, sequence generation, and bioinformatic analysis is openly available on GitHub at https://github.com/angusgcunningham/plasmidbackbonedesign. The fine-tuned ft15k model checkpoint used in this study is hosted on HuggingFace at https://huggingface.co/ UCL-CSSB/PlasmidGPT-SFT. The complete training datasets, all generated plasmid sequences, and all processed results required to reproduce the figures and analyses in this manuscript are archived on Zenodo at https://doi. org/10.5281/zenodo.17820200. These repositories together provide full transparency and enable complete reproducibility of the computational and experimental workflows described in this work.

---

**Supplementary Figure S1:**
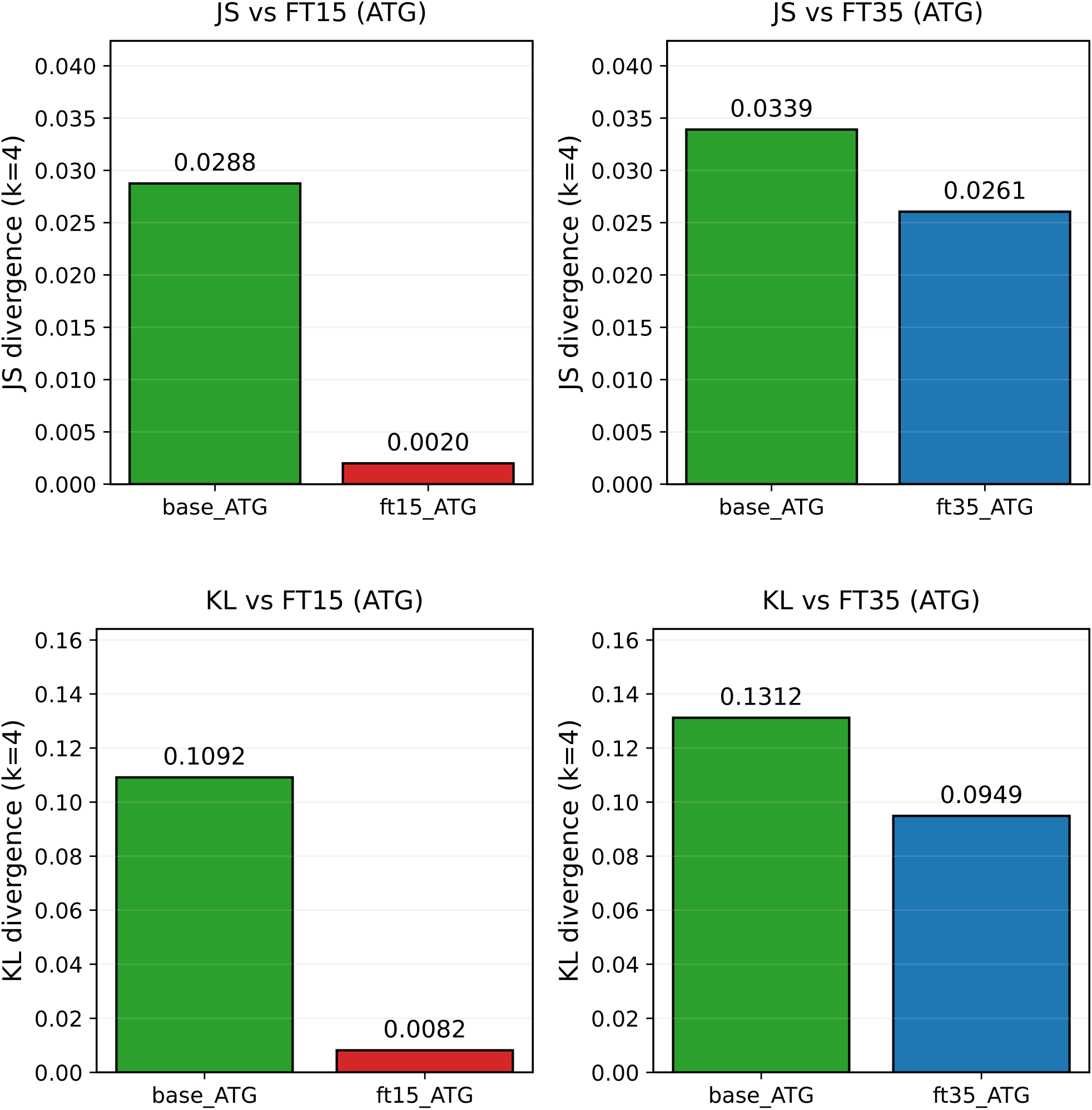
Side-by-side comparison of 4-mer JS divergence (Top) and KL divergence (Bottom) for ATG-seeded models. Left: base vs ft15 ATG generations relative to the ft15k dataset set. Right: Base vs ft35 ATG generations relative to the ft35k dataset.

**Supplementary Figure S2:**
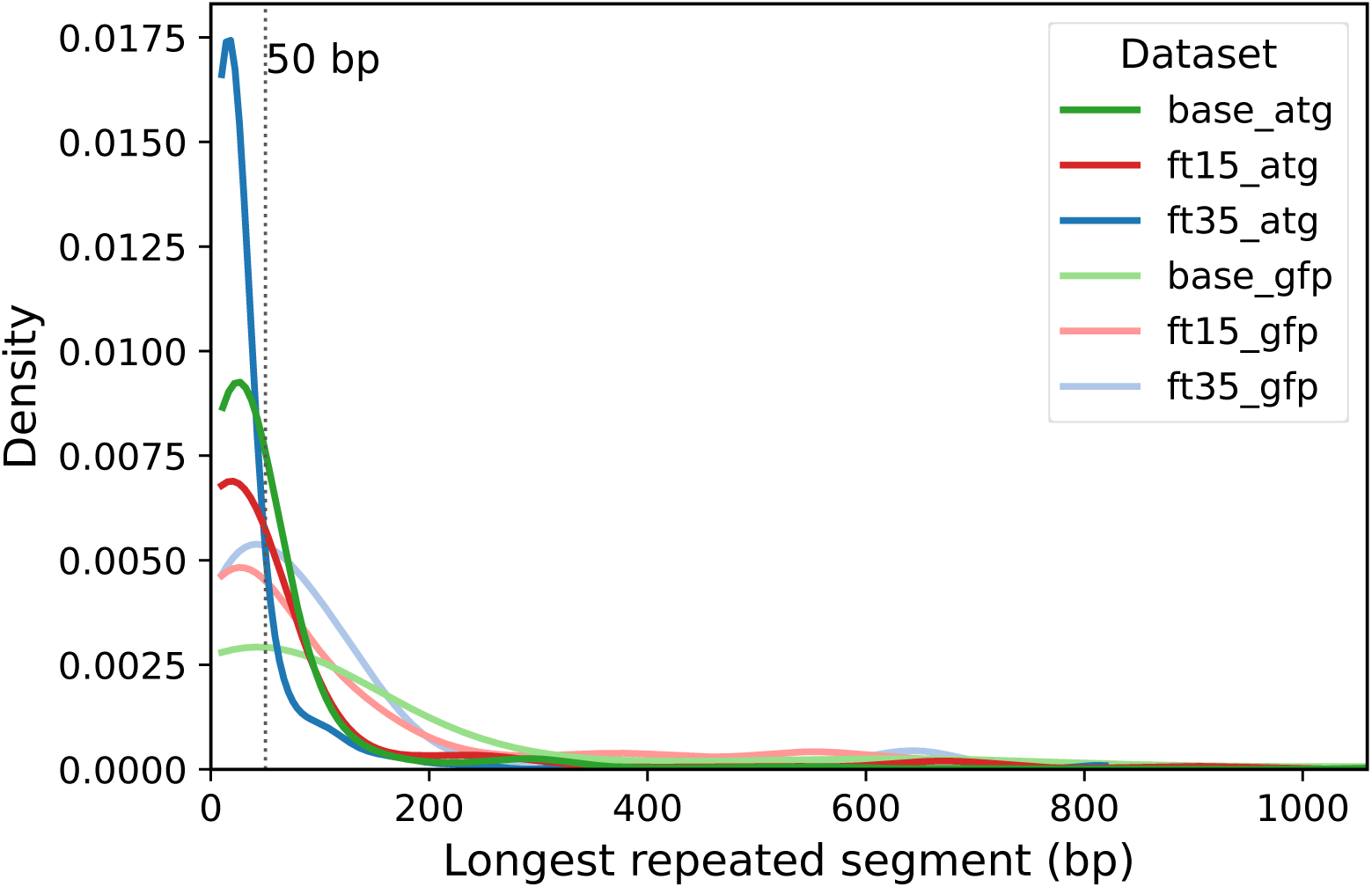
Distribution of repeat lengths. Density plots of the longest repeated segment detected per plasmid across all datasets. A vertical reference line at 50 bp indicates the threshold used in downstream filtering.

**Supplementary Figure S3:**
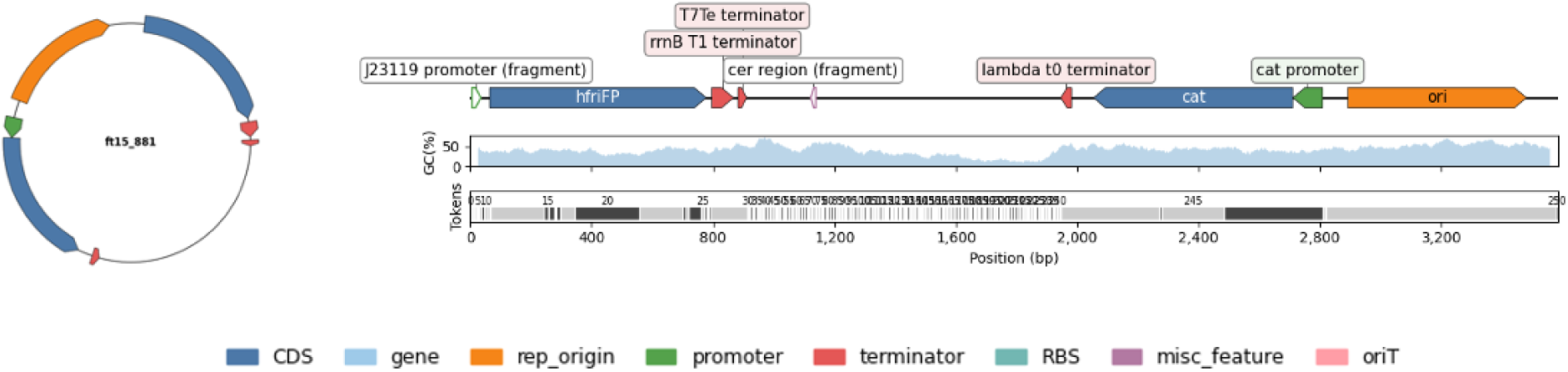
Realisation of failed plasmid design. Representative circular and linear backbone maps for synthesis candidate ft15 891. Failed due to low GC window despite passing strict filtering. GC content(%) and token alignment displayed.

**Supplementary Figure S4:**
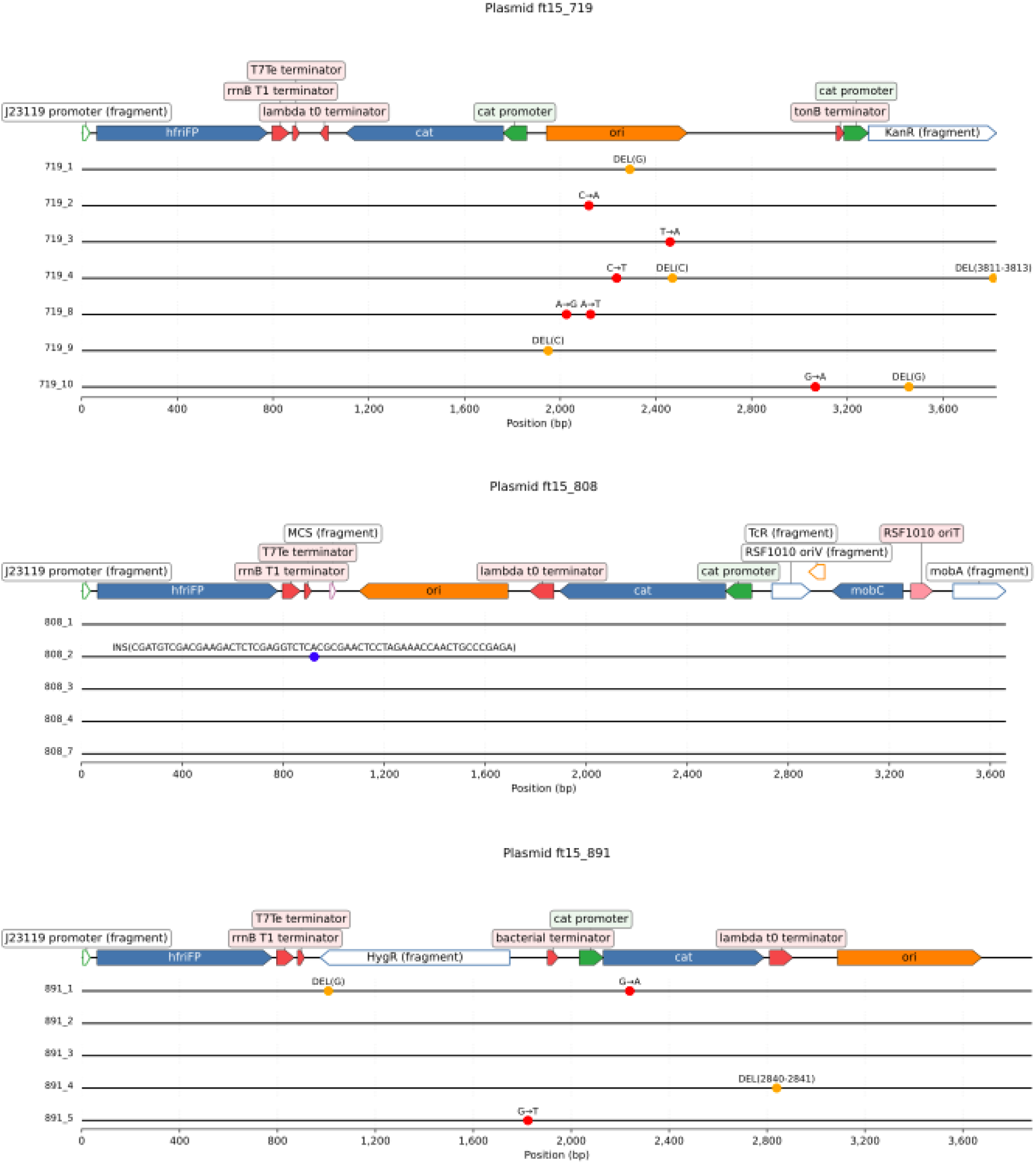
Alignments of transformed plasmid colonies. Representative linear backbone maps from synthesis candidate backbones and their colonies from wet-lab transformations(ft15 719, ft15 808, ft15 891). The aligned mutations represent the position on the backbone of deletions (yellow), substitutions (red) and insertions (blue).

**Supplementary Table S1:** A full list of chemicals and reagents used in this study.

| Item | Supplier | Product number |
| --- | --- | --- |
| Agar | Millipore | 5039 |
| Ampicillin Sodium Salt | BioChemica | A0839 |
| BbsI-HF | New England Biolabs | R3539S |
| Breathe-easy Sealing Membrane | Electron Microscopy Sciences | 70536-10 |
| BsaI-HF v2 | New England Biolabs | R3733L |
| Calcium chloride – dihydrate | North England Laboratory Supplies | 22317.26 |
| Casamino Acids | Millipore | 2240-OP |
| Chloramphenicol | MP Biomedicals Inc | 0219032125 |
| DpnI | New England Biolabs | R0176L |
| Glycerol | VWR | 24388.295 |
| IPTG | Sigma Aldrich | I6758 |
| Kanamycin Sulfate | Fluorochem | M02038 |
| LB Broth | Sigma Aldrich | L3022 |
| M9 Minimal Salts (5x) | Sigma Aldrich | M9956 |
| Magnesium sulfate | Sigma Aldrich | M7506 |
| Microplate 96-Well PS μClear Black Med. | Greiner Bio-One | 655096 |
| Monarch<sup>®</sup> Spin PCR & DNA Cleanup Kit | New England Biolabs | T1130S |
| Monarch<sup>®</sup> Plasmid Miniprep Kit | New England Biolabs | T1010L |
| NEB<sup>®</sup> 5- Competent *E. coli* (High Efficiency) | New England Biolabs | C2987U |
| NEBuilder<sup>®</sup> HiFi DNA Assembly Master Mix | New England Biolabs | E2621L |
| Q5<sup>®</sup> High-Fidelity 2x Master Mix | New England Biolabs | M0492L |
| T4 DNA Ligase (HC) | Promega | M179A |
| T4 DNA Ligase Buffer (10X) | Promega | C126B |
| Thiamine Hydrochloride | Sigma Aldrich | T4625 |
| X-gal | Sigma Aldrich | 9660 |

**Supplementary Table S2:** Strains and plasmids used within this study (RBS = ribosome binding site, CDS = coding DNA sequence).

| Name | Description | Source |
| --- | --- | --- |
| **Strains** |  |  |
| *E. coli* NEB<sup>®</sup> 5-α | Commercial cell line used for cloning | New England Biolabs |
| **Plasmids** |  |  |
| 719 | 719 backbone, J23106 promoter, B0034m RBS, E0040m CDS, B0015 terminator (CmR) | This study |
| 808 | 808 backbone, J23106 promoter, B0034m RBS, E0040m CDS, B0015 terminator (CmR) | This study |
| 891 | 891 backbone, J23106 promoter, B0034m RBS, E0040m CDS, B0015 terminator (CmR) | This study |
| GFP (positive control) | DVC261_AE backbone, J23106 promoter, B0034m RBS, E0040m CDS, B0015 terminator (CmR) | This study |
| DVC261_AE (negative control) | MoClo compatible backbone (CmR) | Casey Chen |

**Supplementary Table S3:**
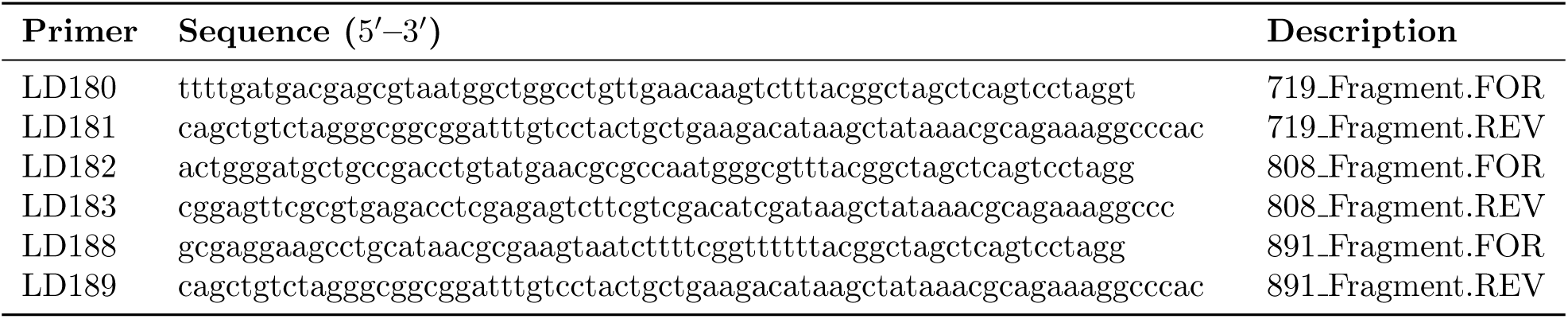
Primer sequences used in this study. All sequences are written 5^′^–3^′^.

